# Autofluorescence imaging of 3D tumor-macrophage microscale cultures resolves spatial and temporal dynamics of macrophage metabolism

**DOI:** 10.1101/2020.03.12.989301

**Authors:** Tiffany M. Heaster, Mouhita Humayun, Jiaquan Yu, David J. Beebe, Melissa C. Skala

## Abstract

Macrophages within the tumor microenvironment (TME) exhibit a spectrum of pro-tumor and anti-tumor functions, yet it is unclear how the TME regulates this macrophage heterogeneity. Standard methods to measure macrophage heterogeneity require destructive processing, limiting spatiotemporal studies of function within the live, intact 3D TME. Here, we demonstrate two-photon autofluorescence imaging of NAD(P)H and FAD to non-destructively resolve spatiotemporal metabolic heterogeneity of individual macrophages within 3D microscale TME models. Fluorescence lifetimes and intensities of NAD(P)H and FAD were acquired at 24, 48, and 72 hours post-stimulation for mouse macrophages (RAW 264.7) stimulated with IFN-γ or IL-4 plus IL-13 in 2D culture, validating that autofluorescence measurements capture known metabolic phenotypes. To quantify metabolic dynamics of macrophages within the TME, mouse macrophages or human monocytes (RAW264.7 or THP-1) were cultured alone or with breast cancer cells (mouse PyVMT or primary human IDC) in 3D microfluidic platforms. Human monocytes and mouse macrophages in tumor co-cultures exhibited significantly different FAD mean lifetimes and greater migration than mono-cultures at 24, 48, and 72 hours post-seeding. In co-cultures with primary human cancer cells, actively-migrating monocyte-derived macrophages had greater redox ratios (NAD(P)H/FAD intensity) compared to passively-migrating monocytes at 24 and 48 hours post-seeding, reflecting metabolic heterogeneity in this sub-population of monocytes. Genetic analyses further confirmed this metabolic heterogeneity. These results establish label-free autofluorescence imaging to quantify dynamic metabolism, polarization, and migration of macrophages at single-cell resolution within 3D microscale models. This combined culture and imaging system provides unique insights into spatiotemporal tumor-immune crosstalk within the 3D TME.

## INTRODUCTION

Immune cells are a critical component of the tumor microenvironment (TME) and regulate tumor initiation, progression, and therapeutic response^1^. The phenotype and function of immune cells surrounding or infiltrating the tumor are crucial to tumor fate^1–3^. Macrophages are a subset of immune cells with diverse function that are abundant within numerous tumor types, especially breast and pancreatic tumors^1,2,4^. This prevalence suggests that macrophages may serve as important immune mediators of tumor behavior, yet macrophage activity is complex and not well understood. For example, multiple macrophage phenotypes have been identified, each with unique functional and stimulatory activity that influence the surrounding cellular environment^5–7^. Two well-characterized classes of macrophages, M1-like and M2-like macrophages, exhibit distinct responses to tumors^7,8^. M1-like macrophages promote anti-tumor activity through apoptotic or phagocytic signaling, secondary immune stimulation via cytokine secretion and antigen presentation, and nutrient deprivation of tumors^7,9,10^. M2-like macrophages support tumor growth by suppressing immune recognition and promoting cancer cell proliferation, motility, and vessel growth^7,9,10^. However, intermediate macrophage states have also been observed within tumors, suggesting a more complex model of macrophage heterogeneity than this M1-M2 dichotomy^11,12^. This heterogeneity within macrophage populations presents a substantial challenge for effective treatments, emphasizing the need to characterize cell-level macrophage function within the 3D TME.

Cell function has been correlated with metabolism across numerous cell types, including macrophages^13^. Therefore, monitoring metabolic preferences within macrophages could reveal heterogeneity in macrophage phenotype and function. Metabolic demands greatly differ between M1-like and M2-like macrophage classes^7,9^. M1-like macrophages generate nitrogen oxide to induce tumor cell death and recruit additional monocytes to the tumor. This increases the production of the metabolic co-enzyme, NADH, via glycolysis and sustains the viability of M1-like macrophages within tumors^14–16^. Conversely, elevated fatty acid oxidation and oxidative phosphorylation in M2-like macrophages support angiogenesis and tumor growth^14–19^. Here, fatty acid oxidation generates acetyl-CoA and NADH, driving downstream oxidative phosphorylation^17,19^. Microenvironmental changes can regulate these distinct metabolic demands and produce diverse macrophage phenotypes^20,21^. This highlights a continuum of metabolic activity related to macrophage function that is provoked by complex stimuli from the TME^20–22^. Therefore, monitoring macrophage metabolic heterogeneity improve understanding of tumor-macrophage crosstalk.

3D tumor models provide reliable models of tumor structure and TME conditions that influence immune cell function. The key advantages of microscale models include controllable features that can be easily tuned to simulate the TME and designs enabling various combinations of culture conditions that support high-throughput assessment^23,24^. Combined tumor and immune cell culture within an extracellular matrix (ECM) further enables paracrine signaling between cell layers and protein-protein interactions necessary for cell motility^25,26^. These microscale platforms also provide a simple approach to recapitulate *in vivo* tumor-associated environmental conditions (e.g., hypoxia, nutrient availability) that regulate both macrophage phenotype and function^23–25,27^. These representative 3D tumor models provide attractive systems to evaluate macrophage heterogeneity in response to spatiotemporal changes in the microenvironment.

The destructive nature of most standard biological assays highlights the need for non-invasive tools to monitor macrophage heterogeneity within intact samples. Flow cytometry requires single-cell tumor dissociation that destroys the native tumor spatial context, followed by fixation and antibody-labeling that can alter cell function^28^. Standard functional assays (e.g., plate reader-based ELISA) only provide bulk cell measurements that cannot capture cell-level heterogeneity^28^. Whole body imaging (e.g., PET/CT, MRI) can track macrophages in 3D but have poor cellular resolution^29,30^. Therefore, the limitations of standard measurements prohibit monitoring spatiotemporal macrophage heterogeneity within the complex 3D TME. This highlights a need for new non-invasive techniques and *in vitro* models to characterize macrophage heterogeneity.

Autofluorescence imaging has previously quantified cellular heterogeneity *in vivo* and in 3D cultures using label-free two-photon microscopy^31–42^. Specifically, autofluorescence imaging can quantify the endogenous fluorescence of metabolic coenzymes NADH and FAD, which are both used across several cellular metabolic processes^43–47^. NADH and NADPH have overlapping fluorescence properties, and are collectively referred to as NAD(P)H ^48^. Fluorescence intensity measurements can inform on the intra-cellular concentrations of NAD(P)H and FAD. The optical redox ratio, defined as the ratio of NAD(P)H intensity to FAD intensity, provides a measure of the relative oxidation-reduction state of the cell^32,35,44^. Fluorescence lifetime imaging microscopy (FLIM) of NAD(P)H and FAD provide additional information specific to protein binding activity^49^. Lifetime measurements can distinguish between free and protein-bound forms of NAD(P)H and FAD, characterized by distinct molecular conformations that affect fluorescence quenching^49^. Previous studies have shown that metabolic autofluorescence imaging detects spatial and temporal changes in stromal cells across *in vivo* and 3D *in vitro* models^31,38,39,42,50–55^. Thus, metabolic autofluorescence imaging of microscale 3D models was demonstrated to quantify metabolic activity and visualize macrophage heterogeneity within the 3D TME using primary human cancer, human cell lines, and mouse cell lines

## METHODS

### 2D cell culture

RAW 264.7 murine macrophages (ATCC TIB-71) were maintained in culture medium composed from standard RPMI 1640 (Gibco), 10% fetal bovine serum, and 1% penicillin:streptomycin. RAW macrophages were seeded in 35 mm glass bottom dishes (MatTek) for imaging experiments. All imaging samples were plated at a density of 1×10^5^ cells per dish and incubated at 37 °C and 5% CO2, for 24 hours to allow cell adhesion. Separate cultures of macrophages polarized with M1-like (IFN-γ) or M2-like (IL-4/IL-13) cytokines were generated by standard media substitution with 2 mL cytokine-supplemented media and incubated between 24-72 hours. Media for M(IFN-γ) stimulation consisted of RPMI 1640 supplemented with 10 ng/ml interferon-γ (IFN-γ; R&D systems), and media for M(IL4/IL13) stimulation consisted of RPMI 1640 with 20 ng/ml interleukin 4 (IL-4; Invitrogen) and 20 ng/ml interleukin 13 (IL-13; Gibco).

### Microdevice design and fabrication

Description of the Stacks microfluidic platform is previously described in detail^27^. 24-well polystyrene microdevices (dimensions: 75 mm × 50 mm) were fabricated by injection molding and sterilized via sonication in isopropanol. Collagen hydrogels (2.0 mg/ml) were prepared and a 4.5 µL volume was suspended within each microwell of the device (thickness ∼1.2 mm), creating an open-air culture system compatible for two-photon imaging. The hydrogel was generated from a mixture of 6 µL 10X PBS, 14 µL sterile water, 6 µL NaOH (Sigma), 160 µL Bovine collagen type I (PureCol, Advanced BioMatrix), 3 µL fibronectin (Sigma) and 50 µL cell type-specific medium (RPMI1640 – mouse; complete DMEM/F12 (Gibco) – human). Collagen hydrogels were polymerized by incubation at 37°C for at least 6 hours, prior to sequential seeding of breast carcinoma cells and macrophages (∼1000 cells/μL each) on the opposing ends of the collagen layer.

### 3D microdevice culture

The 3D microfluidic platform was used for co-cultures of primary and immortalized macrophages and breast carcinoma of both mouse and human origin. PyVMT (established from FVB MMTV-PyVmT mouse tumors) and MDA-MB-231 (ATCC HTB-26) human breast cancer cells were cultured in medium composed from standard Dulbecco’s Modified Eagle Medium (DMEM, Gibco), 10% fetal bovine serum, and 1% penicillin:streptomycin. RAW 264.7 macrophages (ATCC TIB-71) and THP-1 human monocytes (ATCC TIB-202) were cultured in RPMI 1640 medium supplemented with 10% fetal bovine serum and 1% penicillin:streptomycin. Patient-derived invasive breast carcinoma cells (IDC; 171881-019-R-J1-PDC, Passages 7-12) were obtained upon request from the NCI Patient-Derived Models Repository (NCI PDMR)^56^. Culture of patient-derived cells required specialized media consisting of 1X Advanced DMEM/F12 (Gibco) supplemented with 5% Fetal Bovine Serum, 1.1 µM Hydrocortisone (Sigma), 1.61 nM EGF Recombinant Human Protein (Invitrogen), 0.178 mM Adenine (Sigma), 1% penicillin:streptomycin, 2 mM L-Glutamine (Invitrogen), and 0.01 mM Y-27632 dihydrochloride (Tocris)^56^. Well conditions included macrophages incubated with and without tumor cells over 72 hours. For gene expression analysis, tumor cells and macrophages were cultured on separate Stacks layers to prevent cell type cross-contamination. An additional Stacks layer with wells only containing 10 μL culture medium was placed on top of each Stacks layer with cells. These layers were then overlaid form a four-layer Stacks setup (media layer-macrophage layer-media layer-tumor layer). Media changes were performed daily by aspirating old media from each well and replacing with 10 µL of fresh culture media during incubation between imaging experiments.

### Autofluorescence imaging

FLIM images were acquired with an Ultima two-photon imaging system (Bruker) composed of an ultrafast tunable laser source (Insight DS+, Spectra Physics) coupled to a Nikon Ti-E inverted microscope equipped with time correlated single photon counting electronics (SPC 150, Becker & Hickl GmbH). The ultrafast tunable laser source enabled excitation of NAD(P)H (750 nm) and FAD (890 nm) fluorescence. For 2D macrophage imaging, NAD(P)H and FAD fluorescence were excited sequentially, and their respective emission was isolated using 440/80 and 550/100 bandpass filters (Chroma). Laser power at the sample for NAD(P)H and FAD excitation was approximately 11.3 mW and 11.5 mW, respectively. Simultaneous excitation of NAD(P)H (750 nm) and FAD (895 nm) was performed for all 3D samples by wavelength mixing of 750 nm and 1040 nm laser lines to virtually generate a 895 nm excitation effect as described previously^57^. Approximate laser power at the sample during wavelength mixing was 10.7 mW at 750 nm and mW at 1040nm. NAD(P)H and FAD emission during wavelength mixing were isolated using a 466/40 and a 540/24 nm bandpass filter (Chroma), respectively. All samples were illuminated through a 40X/1.15 NA objective (Nikon), with 2X optical zoom for wavelength mixing to capture fluorescence emission only from pixels with efficient mixing. Emission was detected with GaAsP photomultiplier tubes (Hamamatsu 7422PA-40). Per-pixel lifetime decay curves were collected by scanning the field of view (256×256 pixels; 300µm X 300 µm) over 60 seconds with a 4.8 microsecond pixel dwell time. To prevent photobleaching within the sample, photon count rates were maintained at ∼1-3×10^5^ photons/second. The instrument response function was generated from second harmonic signal of urea crystals excited at 900 nm, with a full width at half maximum of 240 picoseconds. A fluorescence lifetime standard measurement was collected daily by imaging a YG fluorescent bead (Polysciences Inc), measured as 2.1±0.09 nanoseconds consistent with reported lifetime values^58,59^. Autofluorescence images were captured for each 2D polarization condition at 24, 48, and 72 hours post-stimulation, with a minimum of four representative fields of view per sample (∼1000-2000 cells). For 3D cultures, autofluorescence lifetime volumes were acquired across 2-3 wells for each macrophage monoculture and co-culture condition for at least 3 separate devices (i.e., 6 to 9 wells per condition). Image volumes captured images at 3 µm z-steps starting at the macrophage seeding plane and ending at the leading edge of the macrophage layer (∼1000-3000 cells). Z-stack image depths ranged from 24 to 210 µm.

### Image analysis

Per-pixel fluorescence lifetimes of free and protein-bound NAD(P)H and FAD were calculated from fitting fluorescence decays to the following bi-exponential model: 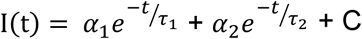 ^49^. Fluorescence intensity images were generated by integrating photon counts over the per-pixel fluorescence decays. The per-pixel ratio of NAD(P)H fluorescence intensity to FAD intensity was calculated to determine optical redox ratio. A customized CellProfiler pipeline was used to segment individual cell cytoplasms^60^. Cytoplasm masks were applied to all images to determine single-cell redox ratio and NAD(P)H and FAD fluorescence lifetime variables. Fluorescence lifetime variables consist of the mean lifetime (τ_m_ = τ_1_α_1_ + τ_2_α_2_), free- and protein-bound lifetime components (τ_1_, τ_2_), and their fractional contributions (α_1_ and α_2_; where α_1_ + α_2_ = 1) for each cell cytoplasm. Nine total variables were analyzed for each cell cytoplasm: NAD(P)H τ_m_, FAD τ_m_, NAD(P)H τ_1_, FAD τ_1_, NAD(P)H τ_2_, FAD τ_2_, NAD(P)H α_1_, FAD α_1_, and optical redox ratio.

### Metabolic Inhibition

To validate that metabolic changes were specific to macrophage polarization, 2D polarized macrophages were treated with a panel of metabolic inhibitors with known activity and imaged post-treatment. RAW264.7 macrophages were stimulated to glycolytic M(IFN-γ) or M(IL4/IL13) phenotype for 72 hours^61–64^. Next, glycolysis was inhibited with media + 10 mM 2-deoxyglucose (2DG; Sigma), FAO was inhibited with media + 100 nM etomoxir (ETO; Sigma), or OXPHOS was inhibited with media + 4mM sodium cyanide (NACN; Sigma)^65–67^. Autofluorescence imaging was performed immediately before adding inhibitor and after treatment for 5 mins (NaCN), 1 hour (2DG), or 24 hours (ETO).

### Immunofluorescence staining and imaging

Following autofluorescence imaging, unstimulated and cytokine-stimulated 2D cultures of RAW264.7 macrophages were stained according to supplier instructions with both PE-conjugated CD86 antibody (Tonbo Biosciences) and FITC-conjugated CD206 antibody (BioRad) to confirm macrophage polarization. Conjugated antibodies were diluted to 1:30 in PBS+goat serum. 2D immunofluorescence images were acquired using the two-photon microscope setup described above. CD206-FITC fluorescence intensity was excited at 890 nm and collected with a 550/100 nm bandpass filter, while CD86-PE fluorescence was excited at 1050 nm and collected with a 650/47 nm filter.

### mRNA Isolation, cDNA Synthesis and real time-quantitative PCR (RT-qPCR)

To assess heterogeneity in macrophage polarization and confirm expected functional phenotypes, expression levels of genes specific to broad macrophage classes (i.e., M1-like, M2-like, mixed phenotypes) were analyzed by qPCR. For 2D cultures, separate cultures of M0, M(IFN-γ), M(IL4/IL13) RAW 264.7 macrophages were maintained in 6-well plates prior to mRNA extraction. For 3D cultures, RAW264.7 macrophages or THP-1 monocytes were cultured alone or with breast cancer cells in the 3D Stacks microfluidic platform. For all experiments, mRNA was extracted at 24, 48 and 72 hours using a Dynabeads™ mRNA DIRECT™ Purification Kit (Thermo Fisher, 61012). Purified mRNA was quantified using a Qubit fluorometer (Thermo Fisher) and a Qubit™ RNA BR Assay Kit (Thermo Fisher, Q10210). mRNA was reverse transcribed using RT^2^ PreAMP cDNA synthesis kit (Qiagen, 330451). The prepared cDNA was preamplified using the RT^2^ PreAMP Primer Mix for Human and Mouse PCR Array (Qiagen, PBH-181Z). cDNA was analyzed by RT-qPCR using a Qiagen RT2 profiler custom panel (Qiagen, PAHS-181Z, CLAM3313, CLAH36077) following supplier instructions. Gene lists for mouse and human experiments are reported in Supplementary Table 2.

### Population density modeling

Population density modeling of single-cell metabolism was used to observe differences in macrophage heterogeneity across 2D polarized cultures and 3D tumor-macrophage cultures over time. Frequency histograms were generated based on single-cell metabolic autofluorescence measurements across samples. Multiple Gaussian probability density functions were fit to each distribution histogram to identify the presence of sub-populations with distinct metabolic activity^37^. Accurate sub-population identification was evaluated by iteratively increasing the number of fitted Gaussian curves and calculating the Akaike Information Criterion (AIC) for each iteration^37,68^. The number of fitted Gaussians yielding the lowest AIC value indicate the optimal fit conditions and number of sub-populations per sample^37,68^.

### Statistics

Tukey’s tests for non-parametric, unpaired comparisons were performed to assess differences in autofluorescence variables between 2D cytokine-stimulated cultures and 3D monoculture and co-culture conditions over 72 hours. Multiple student’s t-tests were performed to assess significance of fold change differences represented as volcano plots for redox ratio changes in 2D macrophage populations in response to metabolic inhibitors and gene expression changes with respect to M0 conditions for 2D mouse macrophages or monoculture for 3D mouse and human macrophages. Coefficient of variation was calculated to evaluate variability across 3D mouse monocultures and co-cultures over 72 hours. The non-parametric, squared-ranks test was performed to assess the equality of coefficients of variation between monoculture and co-culture groups (MATLAB, SquaredRanksTest)^69,70^. Metabolic autofluorescence results are represented as line plots, dot plots, or bar graphs plotted in GraphPad. Heatmaps of autofluorescence variables with respect to depth were generated in MATLAB.

## RESULTS

### Microscale 3D co-culture and metabolic autofluorescence imaging provide innovative and complementary tools to monitor 3D changes in macrophage metabolism within the TME

Autofluorescence imaging can non-destructively monitor live, single-cell metabolism and 3D migration, while 3D microscale co-cultures reliably model tumor complexity with selective environmental control and high-throughput capabilities. Specifically, the Stacks 3D microfluidic system enables multi-cellular cultures with minimal biological material, flexible configuration, and well-characterized environmental gradients^27^. The Stacks system was combined with metabolic autofluorescence imaging to monitor tumor-mediated changes in macrophage metabolism and migration (Fig. 1A)^27^. Microscale 3D co-cultures of tumor cells and macrophages were established in the Stacks 3D system using primary human cancer cells, and cell lines from mouse and human origin (Fig 1A). A collagen-based ECM was polymerized within each well, then monocytes/macrophages were seeded with (co-culture) or without (monoculture) breast cancer cells at the opposing end of the ECM layer (Fig 1A). Both the metabolic autofluorescence and microfluidic culture technologies were validated by imaging macrophages in 2D culture with standard stimulation techniques and in 3D co-cultures of macrophages alone in the Stacks 3D system (Fig. 1B). Finally, the metabolic dynamics of tumor-stimulated macrophages were characterized by collecting NAD(P)H and FAD autofluorescence image volumes of monocytes/macrophages migrating across the ECM at 24, 48, and 72 hours (Fig 1C). Overall, this approach provides unique, quantitative imaging of macrophage spatiotemporal dynamics in the TME that cannot be achieved with conventional assessments that lack single-cell resolution, 3D complexity, and/or non-destructive monitoring capability.

**Figure 1:**
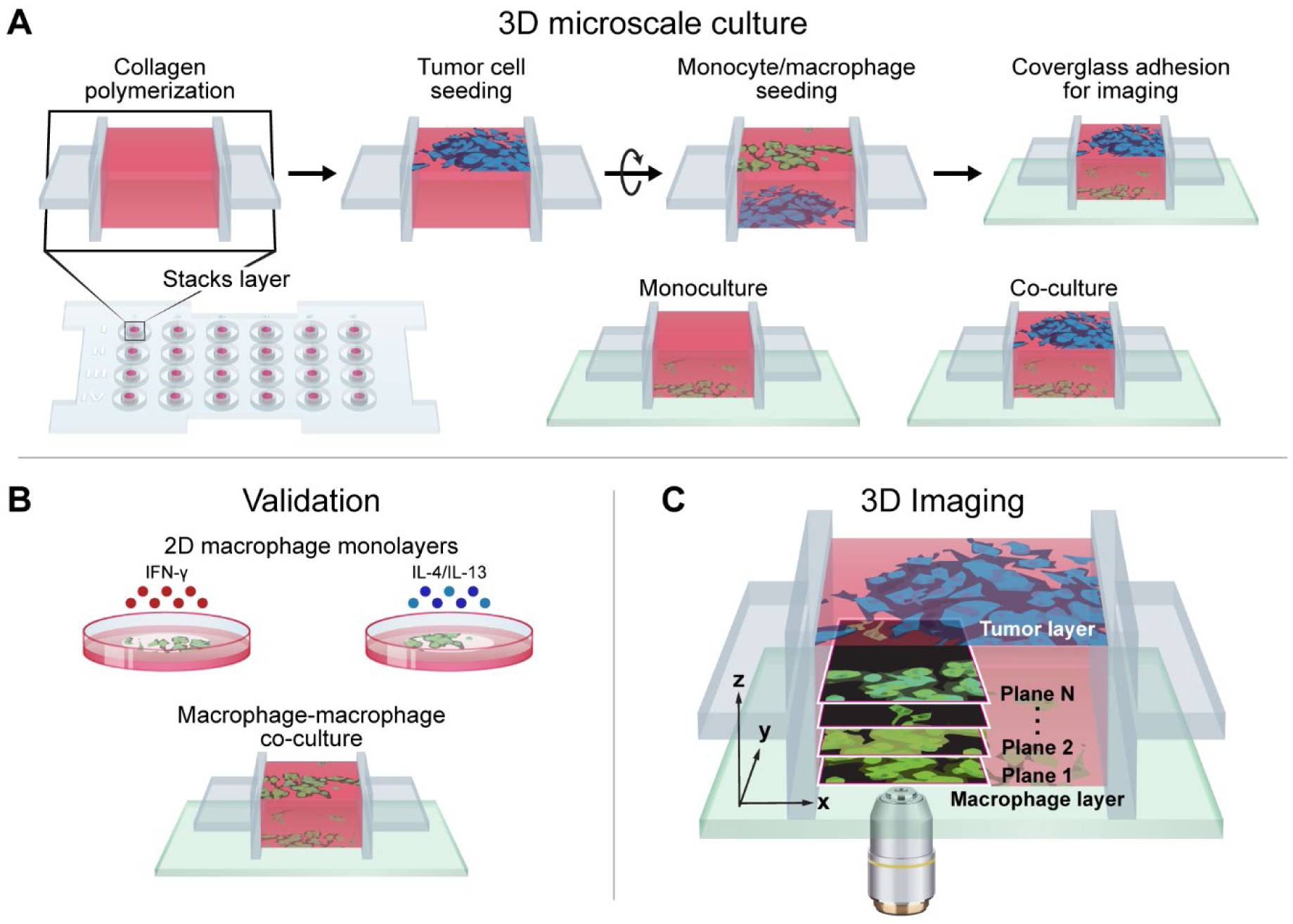
Metabolic autofluorescence imaging of macrophages in the Stacks 3D microscale co-culture system. Illustration of experimental workflow showing (A) design of the Stacks 3D co-culture of macrophages/monocytes and tumor cells from primary patient samples, along with cell lines of mouse and human origin, (B) validation of metabolic autofluorescence imaging in 2D culture with standard stimulations and in Stacks 3D microscale system with macrophages alone, and (C) metabolic autofluorescence imaging of 3D migration and single-cell metabolism for macrophages in the Stacks 3D microscale co-culture system.

### Metabolic imaging validation: macrophage stimulation in 2D *in vitro* culture

To establish the sensitivity of metabolic autofluorescence imaging to distinct macrophage phenotypes, metabolic autofluorescence measurements were first validated in 2D cultures of RAW264.7 mouse macrophages stimulated with IFN-γ (M(IFN-γ): anti-tumor M1-like phenotype), IL-4/IL-13 (M(IL4/IL13): pro-tumor M2-like phenotype), or without cytokine stimulation (M0: naïve macrophages). Macrophages were maintained in cytokine-supplemented media and imaged over 24-72 hours. Autofluorescence imaging of NAD(P)H and FAD demonstrates time-dependent differences in M(IFN-γ) and M(IL4/IL13) macrophages (Fig 2A). M(IFN-γ) macrophages have increased redox ratio compared to M(IL4/IL13) macrophages, with the greatest redox ratio differences at 72 hours post-polarization (Fig 2A). Individual M(IFN-γ) and M(IL4/IL13) macrophages have heterogeneous NAD(P)H and FAD lifetimes across fields-of view at all timepoints. A flattened, cuboid structure was observed for M(IFN-γ) macrophages, whereas M(IL4/IL13) macrophages extended outward with spindle projections, consistent with reported morphology of polarized macrophages (Fig 2A)^71^. These known subset-specific morphological changes become more pronounced over time (Fig 2A). Staining for common M1-like and M2-like surface proteins (CD86 and CD206, respectively) show increasing CD86 expression in M(IFN-γ) macrophages across the 72 hour time course, while M(IL4/IL13) macrophages have stronger CD206 expression (Fig 2A).

**Figure 2:**
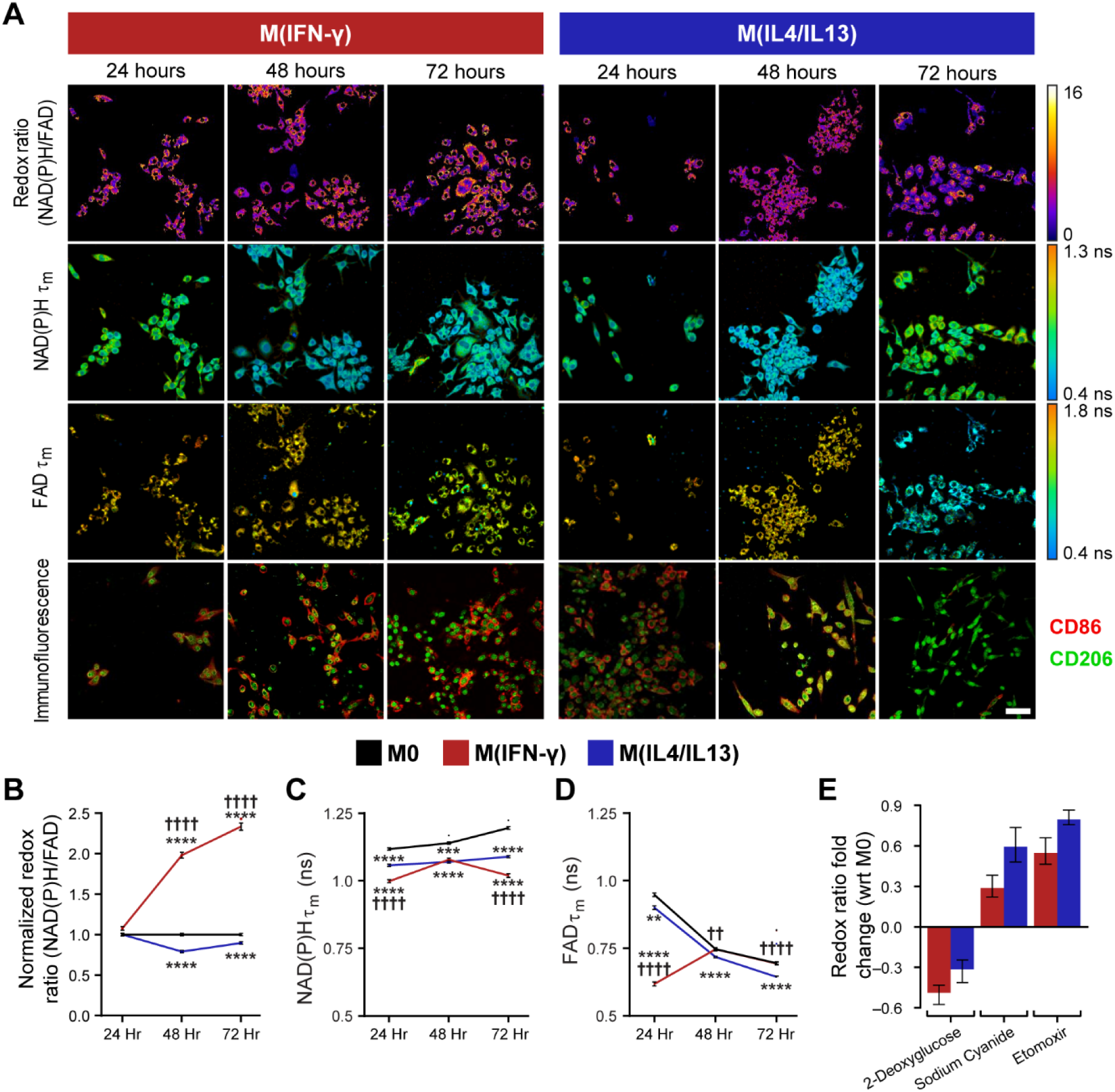
Metabolic autofluorescence imaging is sensitive to temporal changes in macrophage metabolism in 2D culture. A) Representative optical redox ratio, NAD(P)H and FAD mean lifetime (τ_m_) images of RAW 264.7 macrophages cytokine-stimulated to M(IFN-γ) (anti-tumor M1-like phenotype), M(IL4/IL13) (pro-tumor M2-like phenotype), or unstimulated (M0 naïve phenotype). Representative immunofluorescence images show surface expression of known M1-like and M2-like macrophage markers (CD86, CD206). Scale Bar = 50 μm. Quantitative analysis of B) redox ratio and mean lifetimes (τ_m_) of C) NAD(P)H and D) FAD show dynamic metabolic changes during continued exposure to cytokines. **,***, ****p<0.01,0.001, 0.0001 vs. M0; ††, †††† p<0.01,0.0001 vs. M(IL4/IL13). Metabolic characterization was performed via inhibition with E) 2-deoxyglucose (2DG, glycolysis inhibitor), etomoxir (ETO, fatty acid oxidation inhibitor), and sodium cyanide (NaCN, oxidative phosphorylation inhibitor). Reported values are fold change of redox ratio with respect to M0 control under the same polarization conditions.

Quantitative autofluorescence measurements were then compared between M0, M(IFN-γ) and M(IL4/IL13) macrophages over the 72 hour time course to determine whether metabolic autofluorescence can distinguish macrophage polarization states. The redox ratio was significantly different (p<0.0001) between M0, M(IFN-γ) and M(IL4/IL13) macrophages at 48 and 72 hours post-polarization (Fig 2B). M(IFN-γ) macrophages had decreased (p<0.0001) NAD(P)H mean lifetime (τ_m_) at 24 and 72 hours compared with M0 and M(IL4/IL13) macrophages (Fig 2C). However, at 48 hours, M(IFN-γ) and M(IL4/IL13) macrophages had similar NAD(P)H τ_m_ (p>0.05). Significant differences in FAD τ_m_ across all three macrophage conditions were only observed at 24 hours post-stimulation (p<0.05,0.001) (Fig 2D). The short (τ_1_) and long lifetimes (τ_2_) and their relative contribution (α_1_) for NAD(P)H and FAD influence changes in τ_m_ over the time course (Supplementary Fig 1A-F).

Metabolic inhibitors were used to confirm observed autofluorescence differences between M(IFN-γ) and M(IL4/IL13) macrophages. Glycolysis, oxidative phosphorylation, and fatty acid oxidation were inhibited with 2-deoxyglucose, sodium cyanide, and etomoxir, respectively. Fold changes in redox ratio with respect to untreated controls with the same stimulation are shown in Fig. 1E. Inhibition of glycolysis decreased the redox ratio of M(IFN-γ) macrophages to a greater degree than M(IL4/IL13) macrophages (Fig 1E), consistent with published studies that show increased reliance on glycolysis for these anti-tumor, M1-like macrophages compared to pro-tumor, M2-like macrophages^14–16^. Conversely, M(IL4/IL13) macrophages showed greater increases in redox ratio than M(IFN-γ) macrophages with inhibition of oxidative phosphorylation and fatty acid oxidation (Fig 1E). This is consistent with prior studies that show increased reliance on oxidative phosphorylation and fatty acid oxidation in M2-like macrophages compared to M1-like macrophages^14–19^. Fold changes are statistically significant between control and inhibitor-treated conditions across all treatments (Supplementary Table 1). Fold change of both NAD(P)H and FAD mean lifetimes with inhibitors are shown in Supplementary Fig 1. These findings agree with previous studies reporting upregulation of glycolysis in M1-like macrophages, while M2-like macrophages preferentially shift towards a more oxidative metabolic state^5,6,12,14,19^. Collectively, these results demonstrate that metabolic autofluorescence imaging can assess macrophage phenotype and metabolism.

Population density modeling of single-cell redox ratios was performed to determine whether metabolic autofluorescence imaging can resolve macrophage heterogeneity. Population density curves revealed heterogeneous metabolic sub-populations of macrophages across conditions and time points (Fig 3A-C). All stimulation conditions and time points have a narrow population of cells with similarly low redox ratio along with a second population of cells with a broad range of higher redox ratios (Fig 3A-C). These distributions illustrate the heterogeneous metabolic state of macrophages on a single-cell level. Population distributions overlap between M0, M(IFN-γ), and M(IL4/IL13) conditions, which highlights a continuum of metabolic activity between conditions. M(IFN-γ) and M(IL4/IL13) macrophages also exhibited greater variability compared to M0 macrophages at 24 and 72 hours (Supplementary Table 3). Increasing numbers of sub-populations are observed from population distribution analysis of single-cell NAD(P)H and FAD τ_m_ during macrophage stimulation, highlighting substantial heterogeneity in enzyme binding activity (Supplementary Fig 1M-R).

**Figure 3:**
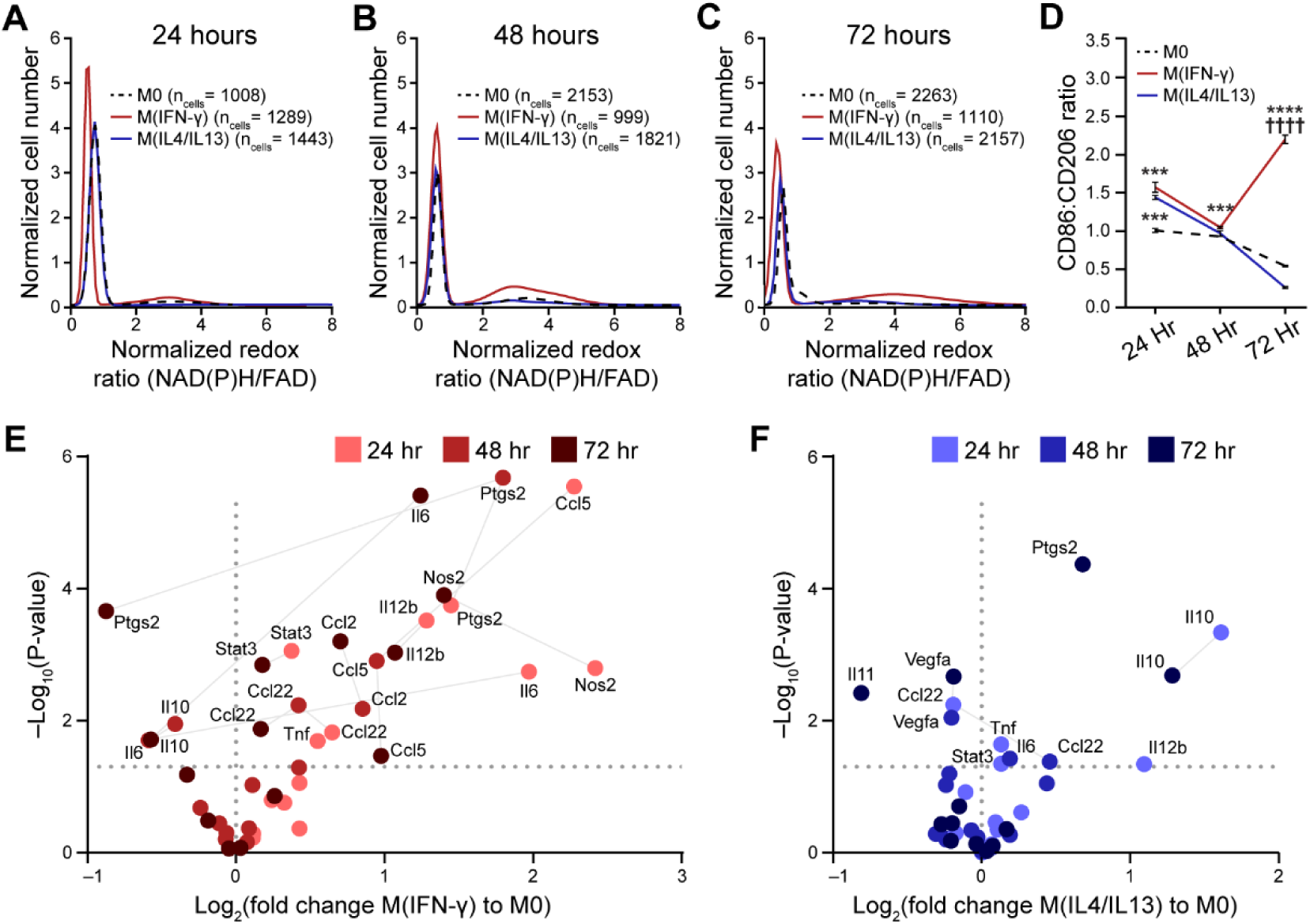
Metabolic autofluorescence measurements resolve metabolic heterogeneity linked to heterogeneous 2D cytokine-stimulated macrophage polarization. Population density modeling of redox ratios per cell illustrates heterogeneous macrophage metabolism at A) 24 hours, B) 48 hours, and C) 72 hours of stimulation. Two distinct sub-populations of cell metabolism are present for each time point and stimulation condition. Expression levels of known M1-like and M2-like macrophage markers (CD86, CD206) were quantified for M0, M(IFN-γ), and M(IL4/IL13) macrophages at D) 24 hours, 48 hours and 72 hours; ** p<0.01, ***p<0.001. Expression levels of M1-like and M2-like genetic markers for E) M(IFN-γ), and F) M(IL4/IL13) conditions were evaluated with qPCR at 24 hours, 48 hours, and 72 hours following stimulation.

Heterogeneity in macrophage function within stimulation conditions (M(IFN-γ) or M(IL4/IL13)) was further assessed from gene and protein expression. Immunofluorescence was quantified from images of M(IFN-γ) and M(IL4/IL13) macrophages stained for CD86 and CD206 expression (Fig 3D). Both stimulation conditions have basal expression of these markers that fluctuate during polarization^72^. Thus, the ratio of CD86 to CD206 expression was used to assess protein changes associated with macrophage polarization^73^. The CD86:CD206 ratio increased significantly in M(IFN-γ) compared with M(IL4/IL13) conditions by 72 hours, while heterogenous expression was observed at earlier time points (Fig 3D). Gene expression of cytokines and other mediators of inflammation (Supplementary Table 2) were also measured with qPCR to assess macrophage function across 24-72 hours of stimulation (Fig 3E-F). Ccl2 and Ccl5 expression and downregulation of IL10 and Ptgs2 in M(IFN-γ) conditions at 72 hours indicates polarization towards M1-like phenotype (Fig 3E). IFN-γ stimulation at 72 hours also upregulated mixed phenotype Il12b, IL6, Ccl22 and Nos2 and M2-like gene, Stat3. Accordingly, IL-4/IL-13 stimulation gradually upregulated IL10 and Ptgs2 while Il1b and Vegfa were significantly downregulated by 72 hours, confirming M2-like polarization (Fig 3F). Variability in expression of M1-like, M2-like, and mixed phenotype genes at 24- and 48-hours of stimulation suggests incomplete polarization at earlier time points (Fig 3E-F).

### Evaluating non-specific effects of 3D Stacks co-culture system on macrophage metabolism and migration

The Stacks system was previously characterized for diffusion gradients and cell viability, survival, and recruitment^27^. To determine non-specific changes in metabolism within the Stacks system, metabolic activity of macrophages 1 hour after seeding was assessed for RAW264.7 macrophage monocultures and RAW264.7+PyVMT mouse breast carcinoma co-cultures. Autofluorescence measurements were not significantly different between monocultured macrophages and co-cultured macrophages, confirming that macrophages were similar for both conditions at the time of seeding (Supplementary Fig 2A-C). Additionally, RAW264.7 macrophages were seeded on both ends of the ECM layer to account for non-specific macrophage migration in the 3D Stacks system without tumor stimulation. Minimal change in macrophage migration was detected over 72 hours within macrophage-macrophage cultures (Supplementary Fig 2D-F). Overall, this data suggests that changes in macrophage migration and metabolism in co-culture at later time-points are due to tumor-specific signaling.

### Mouse macrophage metabolism and migration using 3D metabolic imaging and tumor microscale Stacks system

Next, metabolic autofluorescence was characterized in microfluidic cultures of mouse Polyoma-Middle T Virus (PyVMT) mammary carcinoma and mouse RAW264.7 macrophages over a 72 hour time-course. Redox ratio, NAD(P)H τ_m_ and FAD τ_m_ of co-cultured macrophages from a representative z-plane (migration distance = 6 µm) are visually distinct from monocultured macrophages at the same z-plane as early as 24 hours post-seeding (Fig 4A). Morphological differences also emerge between monocultured and co-cultured macrophages over the 72 hour time course. Co-cultured macrophages were larger than monocultured macrophages and adopted a mixture of flat, cuboidal or elongated, spindle-like morphologies, commonly observed for polarized macrophage states (Fig 4A)^71^. Quantitative autofluorescence measurements more clearly capture metabolic changes across 3D macrophage cultures. Significant differences in NAD(P)H τ_m_ and FAD τ_m_ were observed between monocultured and co-cultured macrophages at 24, 48 and 72 hours, while the redox ratio of co-cultured macrophages significantly decreased only at 72 hours compared to monoculture conditions (Fig 4C-E). The redox ratio gradually decreased over 72 hours of co-culture with tumor cells (Fig 4C), while NAD(P)H and FAD τ_m_ did not significantly change over time (Fig 4D&E). Further changes in NAD(P)H and FAD lifetime components were also observed over the co-culture time course (Supplementary Fig 3A-F).

**Figure 4:**
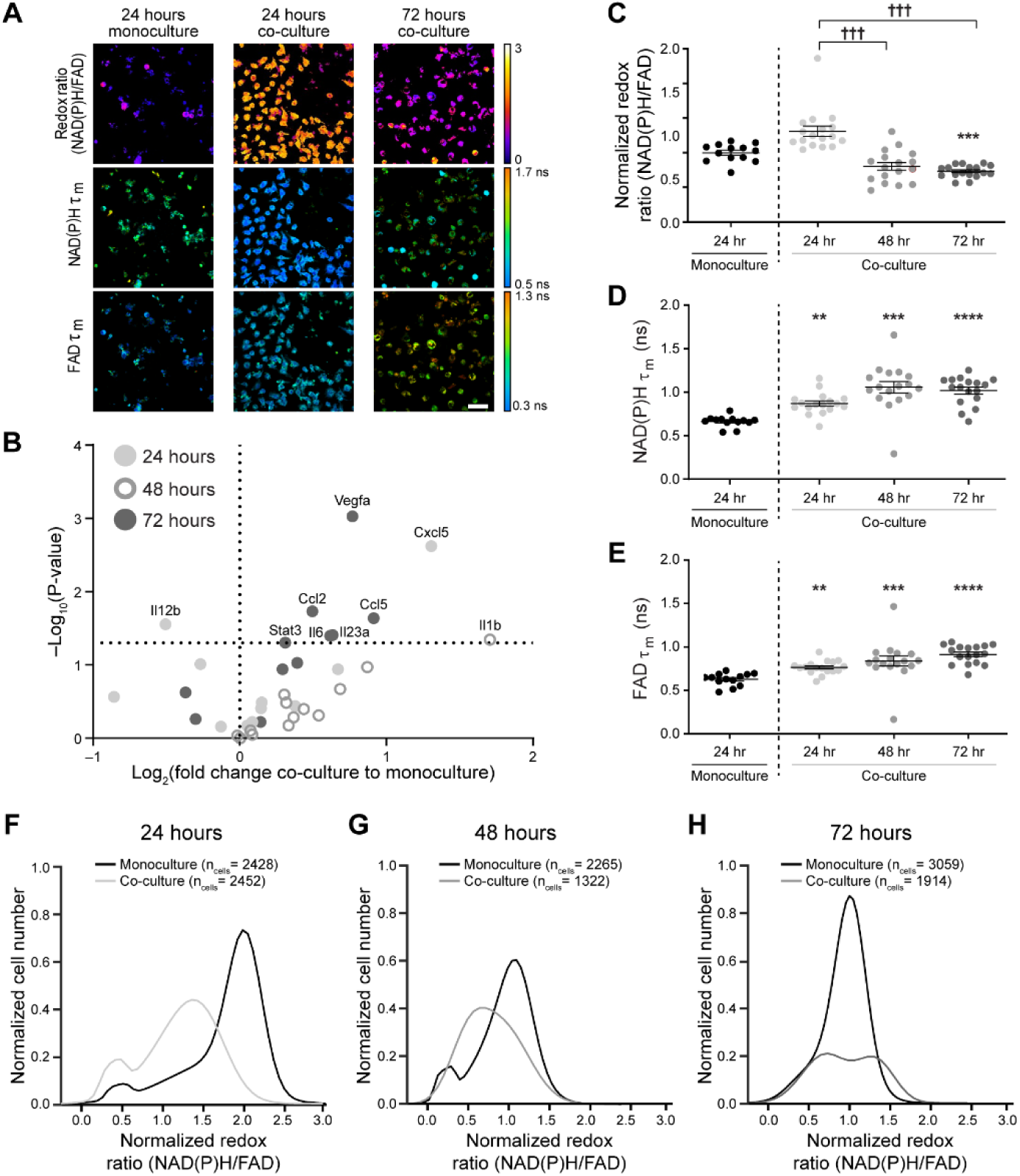
Metabolic autofluorescence imaging of mouse macrophages within 3D Stacks tumor co-cultures. A) Representative autofluorescence images (z-plane distance: 6 µm) demonstrate qualitative differences in the redox ratio, NAD(P)H and FAD mean lifetimes (τ_m_) of RAW264.7 macrophages in co-culture with PyVMT mouse mammary tumor cells. Scale bar = 50 μm. B) Gene expression levels of M1-like and M2-like markers were evaluated for monocultured and co-cultured macrophages at 24, 48, and 72 hours using qPCR. Upregulation of both M1-like and M2-like genes were observed in co-cultures by 72 hours, demonstrating heterogeneity in macrophage polarization in response to tumor stimulation. Quantitative analysis of C) redox ratio, D) NAD(P)H τ_m_, and E) FAD τ_m_ highlight dynamic metabolic changes in macrophages during prolonged tumor-macrophage co-culture. Monocultured macrophages were not significantly different (p>0.05) across time points, so only the 24 hour monoculture is shown above. Significant differences in NAD(P)H and FAD τ_m_ are observed between monocultures and co-cultures, as early as 24 hours post-seeding. Additionally, macrophage co-cultures exhibit a gradual decline in redox ratio over time, consistent with transition to a more oxidized state; **,***,**** p<0.01,0.001,0.001 vs. monoculture, ††† p<0.001. Population density curves of single-cell redox ratios in monocultures and co-cultures at F) 24 hours, G) 48 hours, and H) 72 hours demonstrate variable cellular-level heterogeneity in co-cultured macrophages over time in monocultures.

Genetic heterogeneity between macrophages within 3D co-cultures was evaluated by qPCR for a gene panel associated with M1-like, M2-like, and mixed macrophage phenotypes (Supplementary Table 2)^74^. Minimal changes in gene expression of select M1-like, M2-like, and mixed phenotype genes were observed between monocultured and co-cultured macrophages at early time points (Fig 4B). However, 72 hours of co-culture resulted in significantly increased expression across Ccl2, Ccl5, and IL23a (M1-like), Vegfa and Stat3 (M2-like), and IL6 (mixed phenotype) (Fig 4B). Additionally, Il12b was significantly downregulated in co-cultured macrophages at 72 hours (Fig 4B). Heterogeneity during RAW264.7 macrophage polarization in response to PyVMT tumor cell co-culture was also quantified with population density modeling of cell-level metabolic autofluorescence over the 72-hour imaging time course (Fig 4F-H). Two distinct redox ratio sub-populations of macrophages are present in co-culture at the 24 hour and 72 hour time points, and in monoculture at 24 and 48 hours (Fig 4F&H). Variability in redox ratio of macrophage monocultures is less than co-cultures at all time points (Supplementary Table 3). Heterogeneity in enzyme binding activity is reflected in single-cell NAD(P)H and FAD τ_m_ distributions, which also detect macrophage sub-populations during stimulation (Supplementary Fig 3G-L). These results demonstrate that this 3D tumor co-culture can stimulate heterogeneous changes in macrophage gene expression (Fig. 4B) that may contribute to the metabolic heterogeneity between individual macrophages that is exclusively captured with metabolic autofluorescence measurements (Fig. 4F-H).

Next, mouse macrophage metabolism was monitored during migration toward the tumor layer within 3D co-cultures by acquiring volumes of metabolic autofluorescence images spanning the macrophage layer. Heatmaps of optical redox ratio at each z-plane show that macrophage metabolism and migratory distance vary over the 72 hour time course (Fig 5A). Heatmaps of NAD(P)H τ_m_ and FAD τ_m_ with migration distance are represented in Supplementary Figure 3. The number of macrophages within each z-plane (depth increments of 3 µm) was divided by the total number of macrophages within each image volume to quantify the migration distance of macrophages over time (Fig 5B-D). Monocultured macrophages exhibited minimal migration from the seeding plane, whereas co-cultured macrophages migrated further into the ECM toward the tumor layer especially at 24 and 48 hours (Fig 5B&C). Therefore, the maximum migration distance for the monoculture condition was used to threshold passive from active macrophage migration at each time point (Fig 5B-D)^75^. Actively-migrating macrophages in co-culture have lower redox ratio at all time points, suggesting that these macrophages shift towards oxidative metabolism to enhance migration in the tumor microenvironment (Fig 5E). Additionally, NAD(P)H τ_m_ decreases in actively-migrating macrophages at 24 and 48 hours, while FAD τ_m_ increases significantly at 72 hours (Fig 5F&G). These results highlight the combination of metabolic autofluorescence imaging and 3D Stacks co-culture to reveal novel relationships between metabolic activity and macrophage migration.

**Figure 5:**
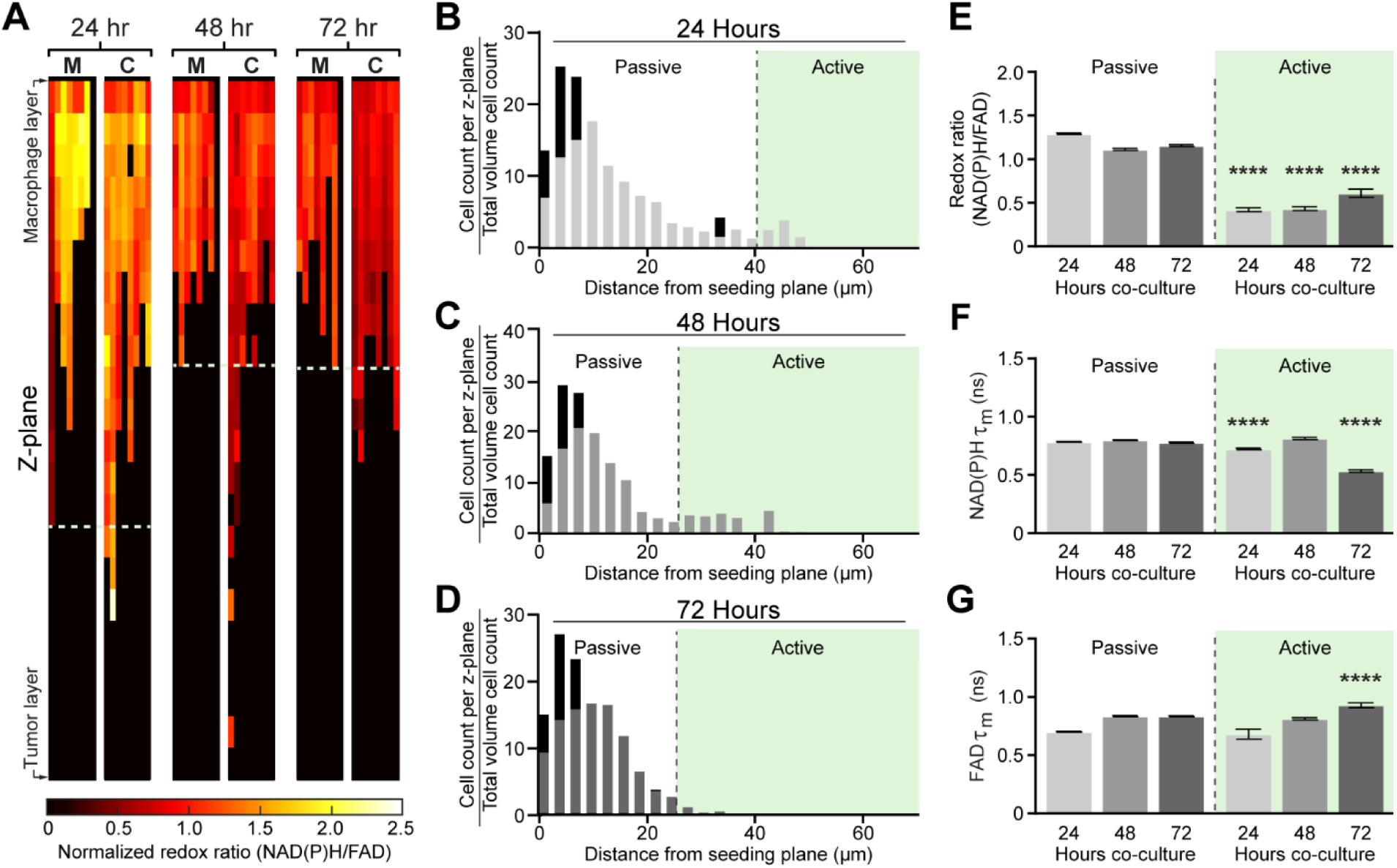
Microscale tumor co-cultures regulate mouse macrophage metabolism and migration over time. A) Representative heatmaps show changes in optical redox ratio with z-plane and time within monocultured, “M”, and co-cultured, “C”, macrophages. Cell density distributions of migration distance identify a distinct subpopulation of actively-migrating RAW264.7 macrophages in the PyVMT mouse mammary tumor co-culture condition at B) 24 hours, C) 48 hours, and D) 72 hours. Passively- and actively- migrating macrophage populations were defined in co-cultures over 72 hours to observe time-dependent relationships between E) redox ratio, F) NAD(P)H τ_m_ and G) FAD τ_m_ and migratory activity. Actively-migrating populations exhibit lower redox ratio than passive-migrating populations at all time points, suggesting macrophages undergo a metabolic switch during migration;****p<0.0001 vs passive migration.

### Human macrophage metabolism and 3D migration in response to patient-derived breast cancer cells

THP-1 human monocytes were cultured alone or with patient-derived IDC cells, and metabolic autofluorescence changes were measured over 72 hours. Qualitative redox images from a representative z-plane (migration distance = 12 µm) highlight cell-level heterogeneity in redox ratio and lifetimes for co-cultures compared to monoculture (Fig 6A). Increased cell and nucleus size is observed in co-cultured cells, consistent with reported morphological differences between human monocytes and activated macrophages (Fig 6A)^76^. Expression of M1-like and M2-like macrophage signature genes demonstrate heterogeneity in macrophage phenotype for human co-cultures ^77,78^. 24 hours of co-culture significantly upregulated only IL1B, while expression of TNF and PTGS2 significantly decreased in 48 hour co-cultures (Fig 6B). Mixed phenotype genes were variably expressed at both 24 and 48 hours (Fig 6B). Conversely, mixed phenotype IL12B, TGFB1, CSF1, NOS2, IL10 was upregulated after 72 hours co-culture with patient-derived tumor cells (Fig 6B). IL1B, TNF (M1-like) and CCL22, VEGFA, PTGS2 (M2-like) were also significantly increased by 72 hours (Fig 6B). Quantitative autofluorescence measurement showed monocyte-derived macrophages in co-culture have significantly higher redox ratio at 48 hours (Fig 6C), no change in NAD(P)H τ_m_ at any time point (Fig 6D), and significantly lower FAD τ_m_ at all time points compared to monoculture (Fig. 6E). NAD(P)H and FAD lifetime components were also affected by tumor co-culture (Supplementary Fig 4A-F). In contrast, the time course of autofluorescence changes in THP-1 human monocytes in co-culture with a triple negative breast cancer cell line (MDA-MB-231, Supplementary Fig 5) substantially differs from patient-derived IDC co-cultures (Fig. 6), highlighting breast cancer cell origin as a source of variability in monocyte-derived macrophage metabolism. These findings demonstrate variation in tumor-mediated monocyte activation and macrophage polarization across species, culture in 2D vs. 3D, and breast tumor sub-types, which highlights the unique insights enabled by combining 3D Stacks and autofluorescence imaging technologies.

**Figure 6:**
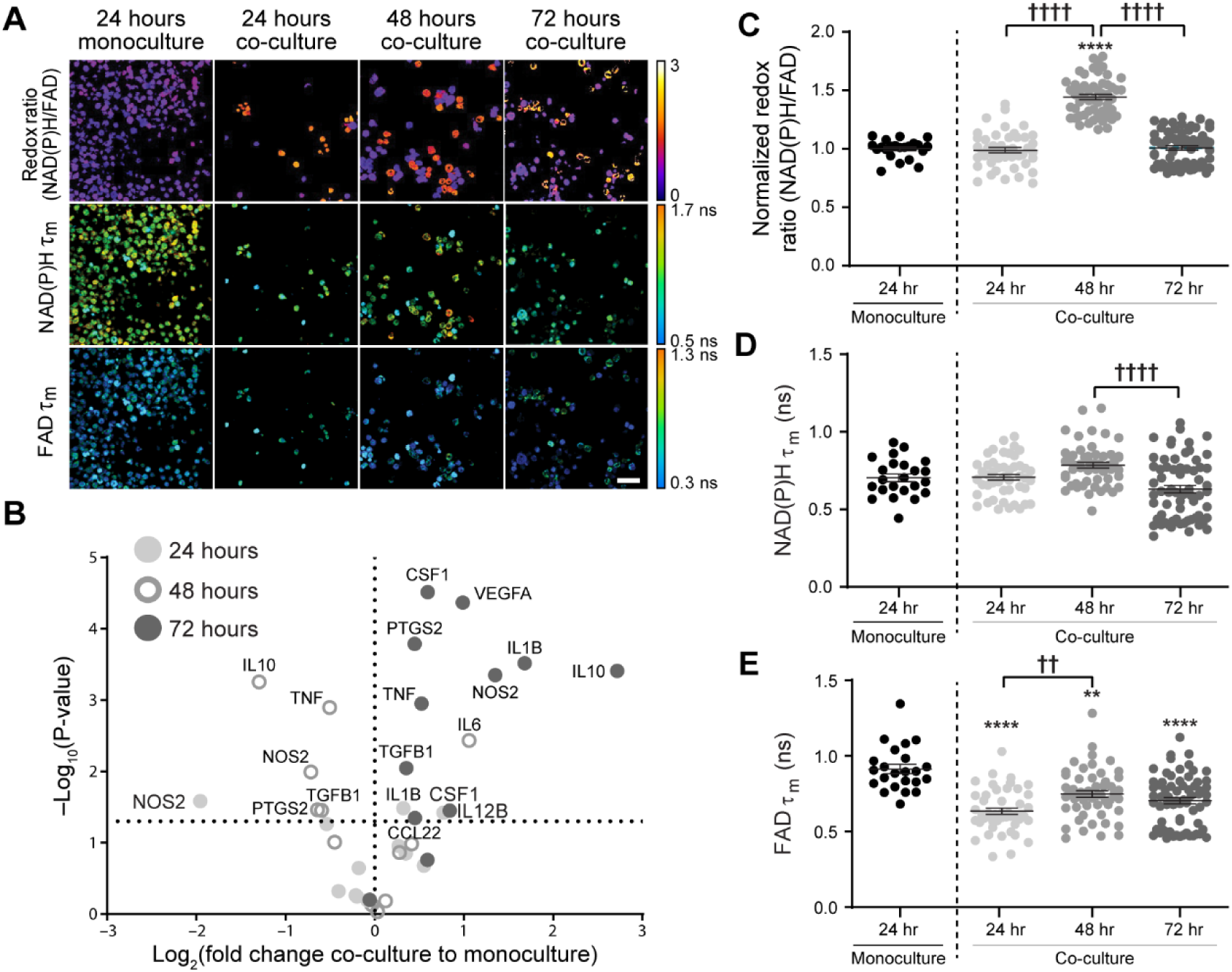
Metabolic autofluorescence imaging captures human monocyte-derived macrophage metabolism in 3D Stacks co-cultures with primary human invasive ductal carcinoma. A) Representative images (z-plane distance: 12µm) display qualitative changes in redox ratio, NAD(P)H τ_m_, and FAD τ_m_ between monoculture and co-cultures over 72 hours. Scale bar = 50 µm. B) Gene expression changes in co-cultured monocyte-derived macrophages over 24, 48, and 72 hours measured from qPCR. Changes are reported as log fold change with respect to the average of the monoculture condition for each measurement and time point. Metabolic changes in human THP-1 monocytes were quantified following co-culture with primary human IDC: C) Redox ratio, D) NAD(P)H τ_m_, and E) FAD τ_m_. **,**** p<0.01,0.0001 vs. monoculture; ††, †††† p<0.01,0.0001.

Tumor-mediated migration was also assessed in 3D co-cultures of primary IDC and THP-1 monocytes. Heatmaps of redox ratio changes with z-plane show increased migration in co-culture conditions and decreased redox ratio in macrophages localized closer to the tumor at 72 hours (Fig 7A), similar to the behavior of RAW264.7 cells (Fig. 5A). NAD(P)H and FAD τ_m_ of macrophages also vary with time and z-plane (Supplementary Fig 4G-H). Monocyte-derived macrophages stimulated in co-culture exhibited substantial migration towards the tumor layer at all time points, in contrast to monocytes in monoculture (Fig 7B-D). Passively- and actively-migrating macrophages were then defined based on monoculture migration distances at each time point (Fig 7B-D). Both the redox ratio and FAD τ_m_ of actively-migrating macrophages co-culture were higher at early time points compared to passively-migrating macrophages, but gradually declined over 72 hours (Fig 7E&G). Actively-migrating macrophages in co-culture also have decreased NAD(P)H τ_m_ at 24 and 72 hours (Fig 7F). Overall, these results establish the versatility of the platform to address the challenge of evaluating dynamic macrophage function in the human TME.

**Figure 7:**
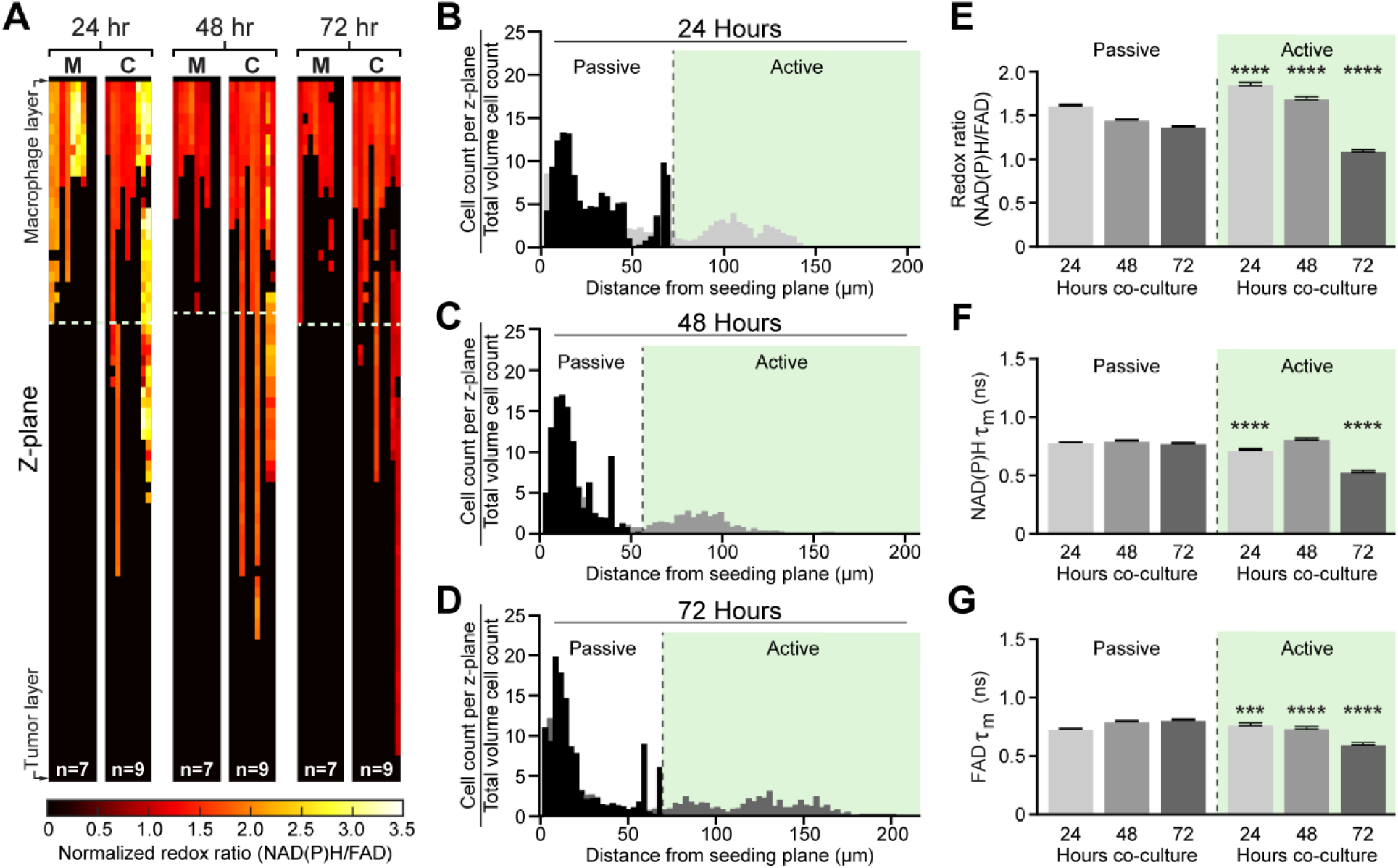
Spatial and temporal metabolic changes of human monocyte-derived macrophages in co-culture with primary human invasive ductal carcinoma. A) Representative heatmaps depicting changes in redox ratio with z-plane. Co-cultured macrophages, “C”, exhibit greater migratory activity and gradual decreases in redox ratio over time compared to monocultured macrophages, “M”. Representative cell density distributions reveal significant monocyte-derived macrophage migration in response to tumor co-culture, with a large population of actively-migrating macrophages at B) 24 hours, C) 48 hours, and D) 72 hours. Differences in E) redox ratio, F) NAD(P)H τ_m_, and G) FAD τ_m_ between actively- and passively-migrating populations were quantified at each time point. ***,**** p<0.001,0.0001 vs. passive migration.

## DISCUSSION

Macrophage plasticity leads to frequent shifts in function and metabolism in the TME, resulting in heterogeneous tumor-associated macrophage populations^14,21,79^. Here, we show that metabolic autofluorescence imaging can monitor macrophage heterogeneity during polarization and migration within a 3D microfluidic model of the TME. Metabolic autofluorescence imaging enables non-destructive monitoring of metabolic changes in individual, live cells within 3D *in vitro* cultures and *in vivo* tumors^31,33,34,39–42,50,51,80^. Additionally, 3D microfluidic culture systems, such as the Stacks microfluidic device used in this study, provide simple platforms to recapitulate dynamic environmental conditions that characterize *in vivo* tumors (e.g., hypoxia, acidosis, nutrient starvation) and are powerful to study primary human cells in an *in vivo*-like environment^81–86^. Previous studies have shown that metabolic autofluorescence can identify macrophages in 2D cultures and *in vivo* but have not explored spatiotemporal heterogeneity in macrophage metabolism within the TME ^34,41,42,53^. This is the first study to monitor cell-level macrophage metabolism in microfluidic models of intact TME. Here, we quantify metabolic heterogeneity of macrophages in the Stacks 3D microdevice platform using autofluorescence imaging to provide novel insights into spatiotemporal heterogeneity of macrophages in the TME.

Temporal changes in microenvironment conditions affect macrophage metabolism and function, so this study quantified time-dependent changes in macrophage autofluorescence within microfluidic 3D tumor co-cultures. This study first confirmed that metabolic autofluorescence imaging can quantify changes in macrophage metabolism and distinguish macrophage sub-populations within intact, living samples. Differences in NAD(P)H and FAD autofluorescence were first observed in 2D cytokine-stimulated macrophages over 72 hours and reflected the expected metabolic shifts for M(IFN-γ) and M(IL4/IL13) macrophages (Fig 2)^87–89^. This result is consistent with previous metabolic flux and metabolite accumulation studies showing increased glycolysis in LPS- or IFN-γ+TNF-α-stimulated (M1-like) macrophages and increased TCA cycle in IL-4-stimulated (M2-like) macrophages, as well as NAD(P)H lifetime studies distinguishing -γ/LPS-treated and IL-4/IL-13 treated mouse bone marrow-derived macrophages (BMDMs)^41^. Next, RAW264.7 macrophages co-cultured in 3D with PyVMT breast cancer displayed changes in redox ratio consistent with metabolic changes from tumor stimulation in previous studies (Fig 4A&C)^90^. Differences in redox ratio and fluorescence lifetime measurements were observed between RAW264.7 macrophages in cytokine-stimulated 2D cultures (Fig 2B-D) and tumor-stimulated 3D cultures (Fig 4C-E), consistent with reported differences in intracellular metabolite concentration and metabolic flux based on stimulation condition (e.g., cytokines, tumor conditioned media) and culture in 2D vs. 3D^91,92^. Additionally, 3D co-cultured RAW264s exhibited heterogeneous cell-level autofluorescence and pooled gene expression (Fig 4B, F-H). This is consistent with reported time courses of metabolic and genetic changes in co-cultures of tumor cells and monocytes or macrophages^93^. Furthermore, the genetic and metabolic heterogeneity observed here is supported by recent studies that similarly show tumor-associated macrophages adopt a mixture of M1-like and M2-like characteristics and functions^94–98^. This reinforces a model of tumor-associated macrophages with diverse phenotypes^99^, which can be further explored using the methods demonstrated here. Ultimately, these studies show that metabolic autofluorescence imaging is sensitive to heterogeneity in macrophage function and metabolism within 3D models of the TME.

Microfluidic culture is also attractive for mimicking the 3D spatial structure of the TME. Accordingly, multi-photon excitation of metabolic autofluorescence enables cellular-resolution imaging in thick, scattering samples to monitor environmental gradients and spatial heterogeneity in 3D tumor models^47,100^. Therefore, we measured metabolic heterogeneity of RAW264 macrophages with 3D migration during co-culture with PyVMT tumor cells (Fig 5). Macrophages in co-culture exhibited greater migration across the ECM layer over 72 hours compared to monocultured macrophages, which was shown to be a tumor-specific behavior (Fig 5A-D, Supplementary Fig 2D-F,3M-N). Previous studies suggest that mouse macrophages often exhibit random migration during chemotaxis^101^. This is consistent with our observations of mouse macrophage migration (Fig 5A-D). Prior studies have also shown that M2-like macrophages are typically more migratory, which is consistent with the oxidative metabolic shift in mouse macrophages localized closest to the tumor layer (actively-migrating) (Fig 5E-G)^102,103^. Overall, metabolic autofluorescence imaging and microfluidic models of 3D microenvironment illustrate tumor-driven spatiotemporal changes in macrophage metabolism.

An important advantage of the Stacks system is the use of primary human cells for studies of human tumor-immune interactions. We highlighted this advantage with 3D co-cultures of primary IDC and THP-1 monocytes (Fig. 6). Previous studies have shown that human THP-1 monocytes will differentiate and preferentially polarize to M2-like macrophages as early as 48 hours after treatment with tumor-conditioned media^104–106^. This is consistent with redox ratio changes of THP-1s in co-culture with primary tumor cells (Fig 6C). The late oxidative, M2-like switch captured by the redox ratio is also consistent with previous reports of 2D differentiated THP-1 cells exposed to IL-4 (M2-like) cytokine stimulation^34^. Co-cultured THP-1s also migrated farther across the ECM than monocultures over 72 hours (Fig 7A-D). Human macrophages and monocytes exhibit strong directional migration toward chemotactic gradients (Fig 7B-D), consistent with our observations of human macrophage and monocyte migration^107^. The redox ratio of actively-migrating human macrophages increased at 24 hours and gradually declined over 72 hours (Figs 7E-G), consistent with reports of upregulated glycolytic metabolism prior to oxidative shifts in human macrophages during chemoattractant-induced migration^108^. These studies indicate that metabolic autofluorescence imaging in human tumor-macrophage microfluidic cultures provides a valuable platform to characterize novel spatiotemporal dynamics of macrophage metabolism within the human TME.

Overall, we have established a novel, single-cell imaging and 3D microfluidic culture platform to monitor the spatial and temporal dynamics of macrophage metabolism within the TME. Autofluorescence imaging of 3D tumor-macrophage co-cultures characterized metabolic heterogeneity with tumor-mediated macrophage polarization and migration. This approach could be used to evaluate metabolic heterogeneity in more complex 3D microscale cultures with additional immune cells, stromal cells, and vasculature. Furthermore, direct manipulation of environmental gradients (e.g., oxygen, nutrients, pH) during culture and imaging could assess the relationship between environmental pressures and macrophage metabolism, function, and organization. Ultimately, these tools may improve our understanding of cellular heterogeneity and metabolism in the TME, and their effects on tumor progression and treatment response.

## ACKNOWLEDGMENTS

We would like to thank Rupsa Datta and Jose Ayuso for their valuable input on the experimental design and manuscript composition. We acknowledge Jens Eickoff for his guidance in statistical analysis of the reported data. We also acknowledge for Matthew Stefely for input on design of scientific diagrams and data representation.

