## Supplemental Figures 1-8 and Supplemental Tables 1-3 for "Autofluorescence imaging of 3D tumor-macrophage microscale cultures resolves spatial and temporal dynamics of macrophage metabolism"

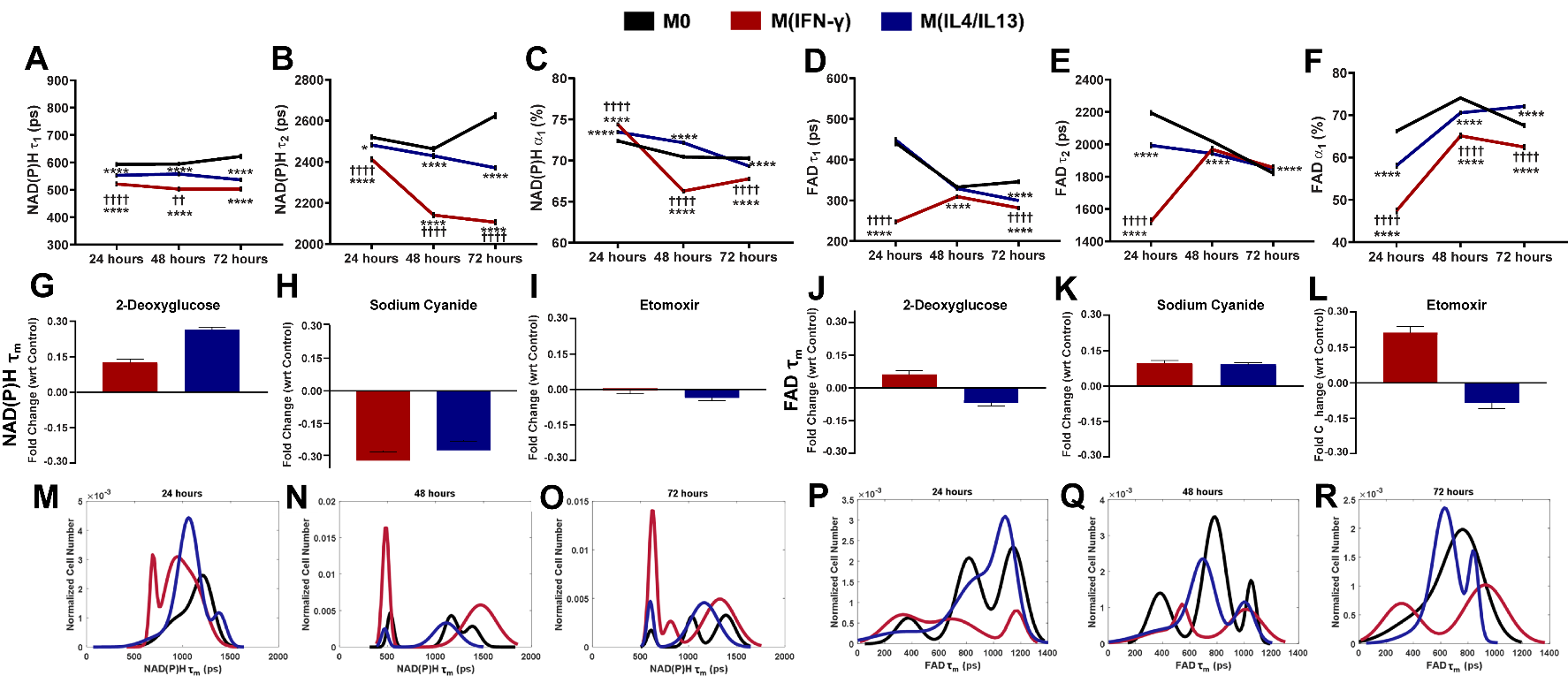

**Supplementary Figure 1: Fluorescence lifetimes of NAD(P)H and FAD exhibit differences between macrophage polarization states.** Quantitative measurement of A) NAD(P)H τ_1_, B) NAD(P)H τ_2_, C) NAD(P)H α_1_, D) FAD τ_1_, E) FAD τ_2_, F) FAD α_1_, illlustrate metabolic differences across 2D cultures of RAW264.7 macrophages polarized to M0, M(IFN-γ), and M(IL4/IL13) over a 72-hour timecourse. Fold change of NAD(P)H τ_m_ in response to treatment with G) 2-deoxyglucose, H) sodium cyanide, and I) etomoxir and FAD τ_m_ in response to treatment with J) 2-deoxyglucose, K) sodium cyanide, and L) etomoxir in M(IFN-γ) and M(IL4/IL13) macrophages shows metabolic inhibitor treatment alters NAD(P)H and FAD mean lifetimes of 2D polarized mouse macrophages. Population distribution modeling demonstrates metabolic heterogeneity for single-cell NAD(P)H τ_m_ over M) 24 hours, N) 48 hours, and O) 72 hours and FAD τ_m_ over P) 24 hours, Q) 48 hours, and R) 72 hours in macrophages unstimulated (M0) and stimulated to M(IFN-γ) and M(IL4/IL13).

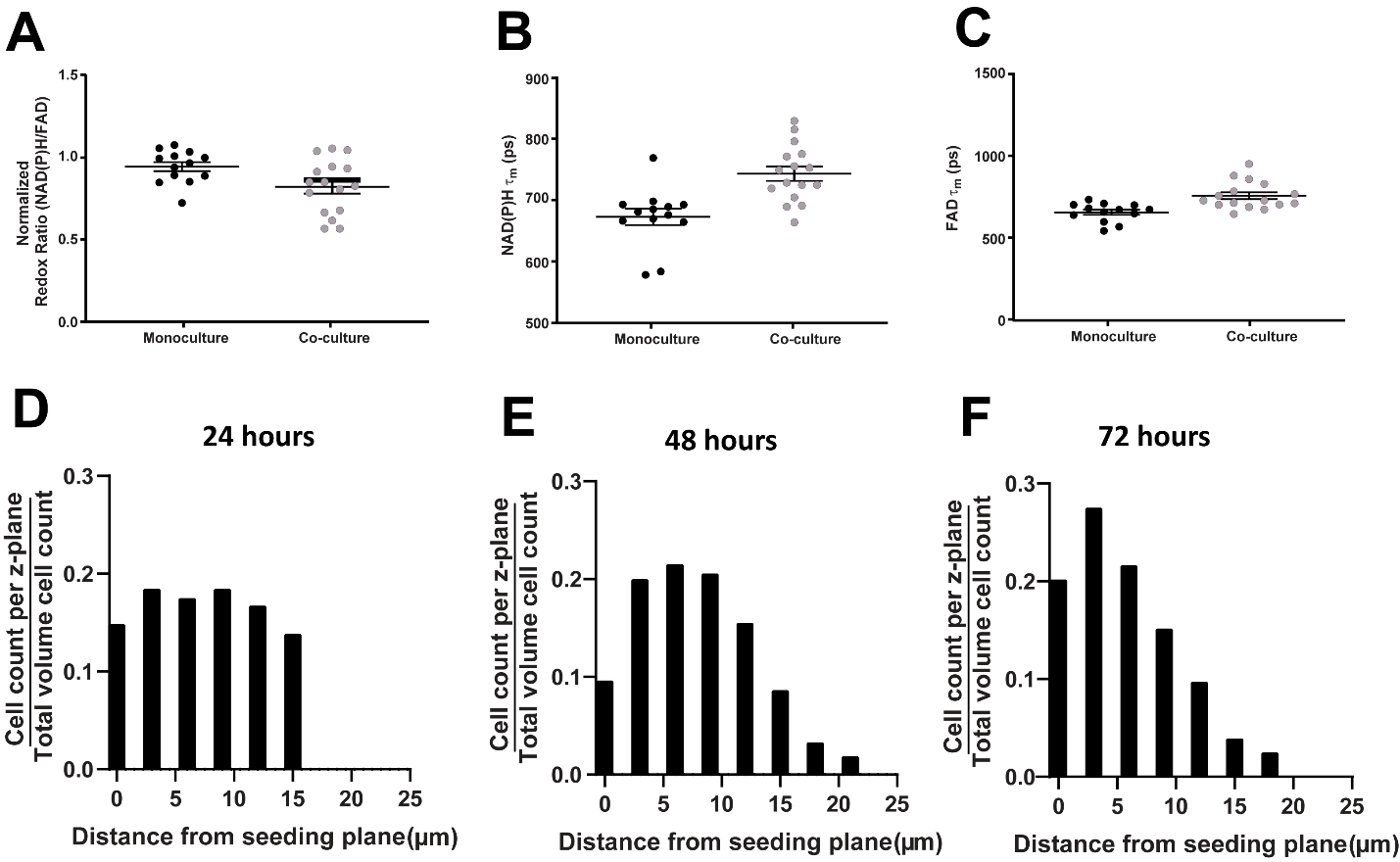

**Supplementary Figure 2: Assessment of non-specific metabolic and migration effects in 3D Stacks co-cultures.** Quantitative measurement of A) redox ratio, B) NAD(P)H τ_m_, and C) FAD τ_m_ across 3D RAW264.7 macrophage monocultures or co-cultures of Polyoma Middle T-virus (PyVMT) breast cancer and RAW264.7 macrophages 1 hour post-seeding. Significant differences were not observed in redox or lifetime measurements between monocultured and co-cultured macrophages, demonstrating metabolic autofluorescence of monoculture and co-cultured moacrophages are similar upon initial seeding. RAW264.7 macrophages were seeded at the top and bottom of 3D ECM layers, and migration was measured at D) 2 hours, E) 48 hours, and F) 72 hours. Migration was quantified from cell counts at each 3 µm slice divided by the total cell count across the entire 3D macrophage layer. RAW264.7+RAW264.7 co-cultures exhibit minimal changes in cell distribution across the collagen layer, suggesting actively-migrating macrophage populations are absent in macrophage-macrophage co-cultures and active migration observed in tumor-macrophage co-cultures is induced by tumor stimuli.

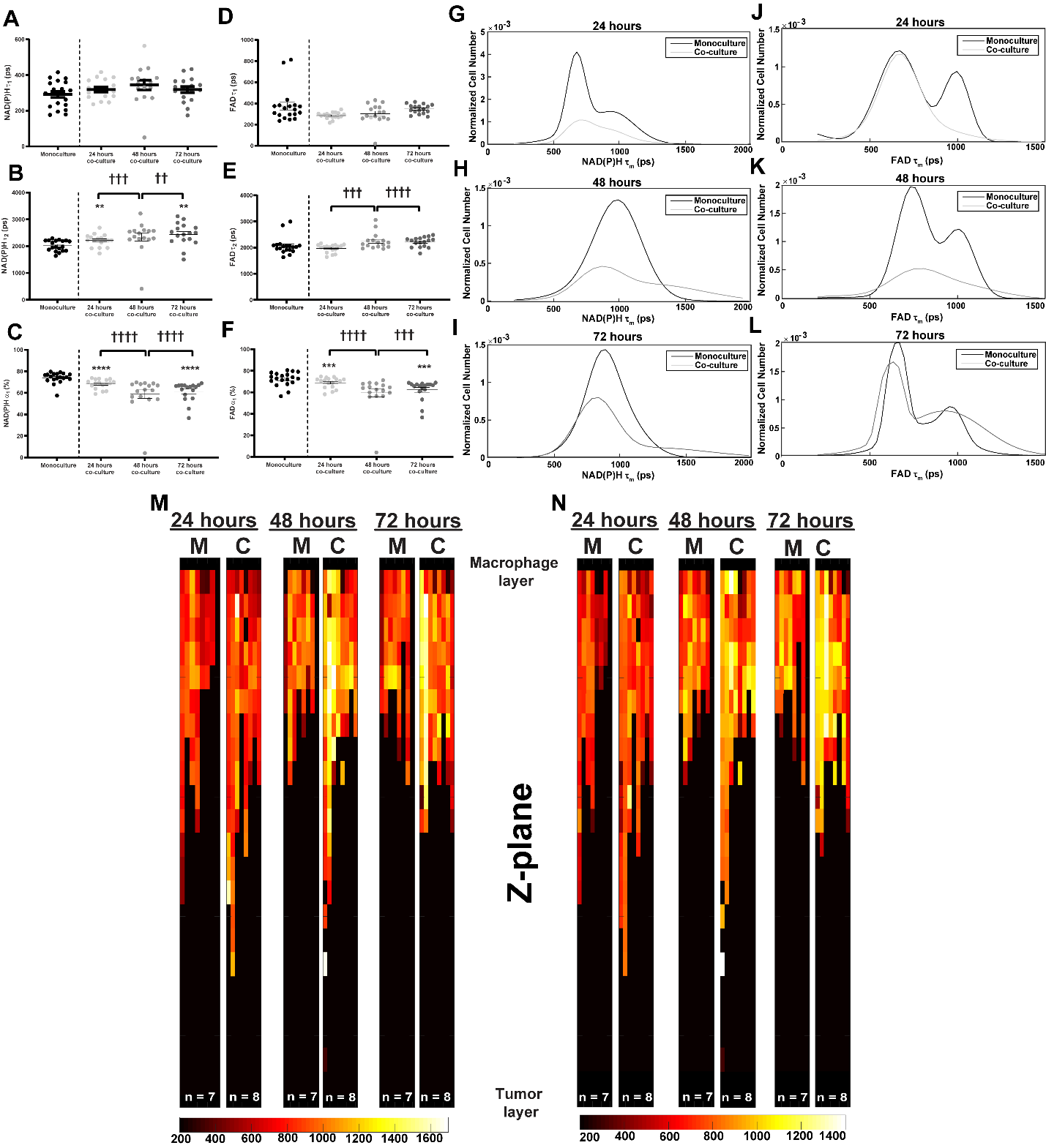
**Supplementary Figure 3: Prolonged co-culture of mouse breast cancer and macrophages yields heterogeneous NAD(P)H and FAD fluorescence lifetime and migration compared to monoculture.** Quantitative measurement of A) NAD(P)H τ_1_, B) NAD(P)H τ_2_, C) NAD(P)H α_1_, D) FAD τ_1_, E) FAD τ_2_, F) FAD α_1_, across 3D RAW264.7 macrophage monocultures or co-cultures of Polyoma Middle T-virus (PyVMT) breast cancer and RAW264.7 macrophages over 24, 48, and 72 hours. Population distribution modeling of single-cell NAD(P)H τ_m_ at A) 24 hours, B) 48 hours, and C) 72 hours post-seeding and FAD τ_m_ at D) 24 hours, E) 48 hours, and F) 72 hours post-seeding in monocultures and co-cultures. Representative heatmaps of M) NAD(P)H τ_m_ and N) FAD τ_m_ during RAW264.7 macrophage migration in 3D monocultures and co-cultures with PyVMT breast carcinoma. Both monocultured and co-cultured macrophages display increased heterogeneity in NAD(P)H and FAD lifetime, regardless of time and migration distance.

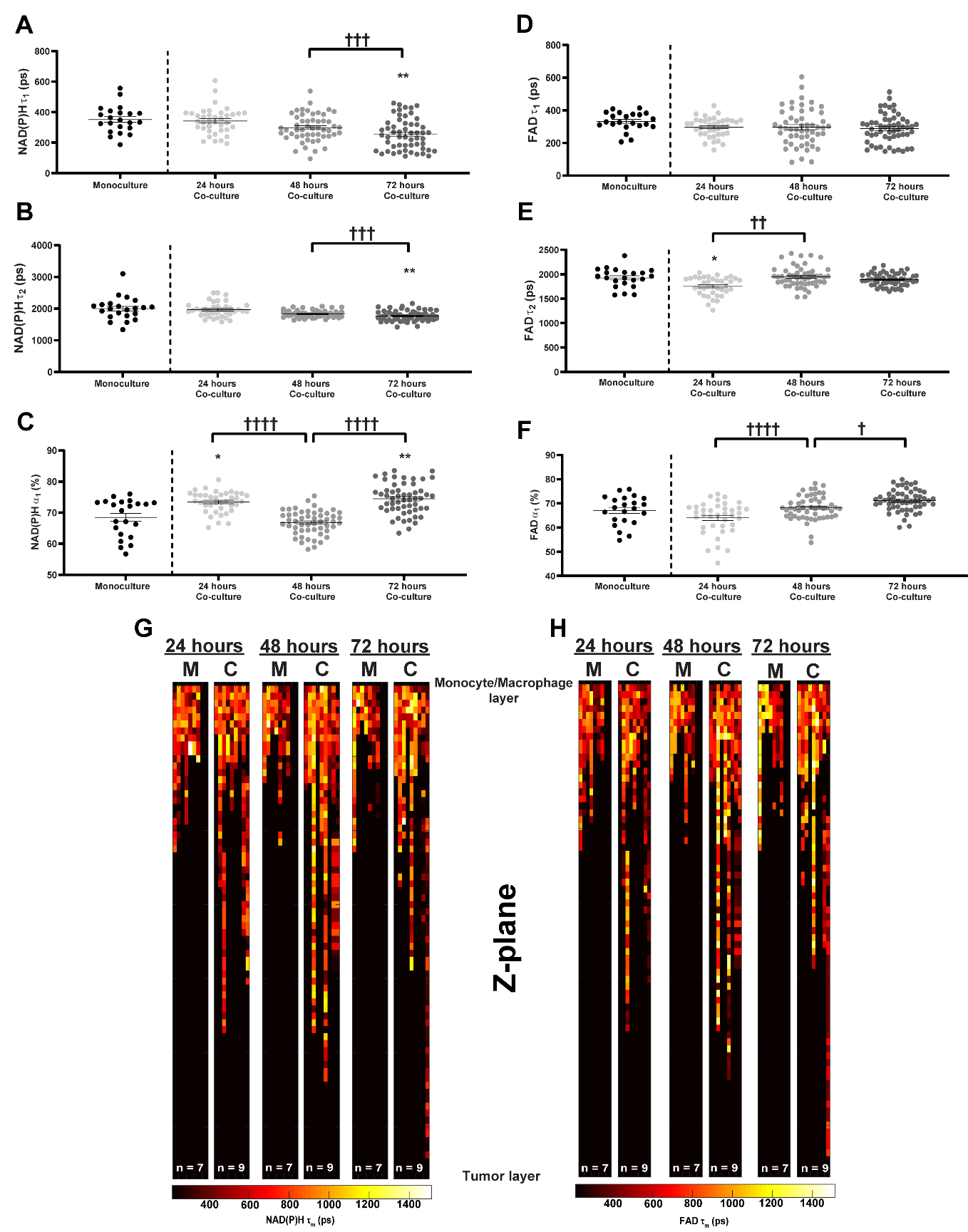

**Supplementary Figure 4: Primary human tumor cells stimulate changes in NAD(P)H and FAD fluorescence lifetime and cell migration in co-cultured human monocyte-derived macrophages.** Quantitative measurement of A) NAD(P)H τ_1_, B) NAD(P)H τ_2_, C) NAD(P)H α_1_, D) FAD τ_1_, E) FAD τ_2_, F) FAD α_1_ across 3D monocultures of human THP-1s or co-cultures of primary breast cancer cells and THP-1s over 24, 48, and 72 hours. Representative heatmaps of G) NAD(P)H τ_m_ and H) FAD τ_m_ during THP-1 migration in 3D monocultures and co-cultures with primary breast carcinoma. Both monocultured monocytes and co-cultured monocyte-derived macrophages exhibit substantial heterogeneity in NAD(P)H and FAD lifetime within z-planes across the 3D layer at all timepoints.

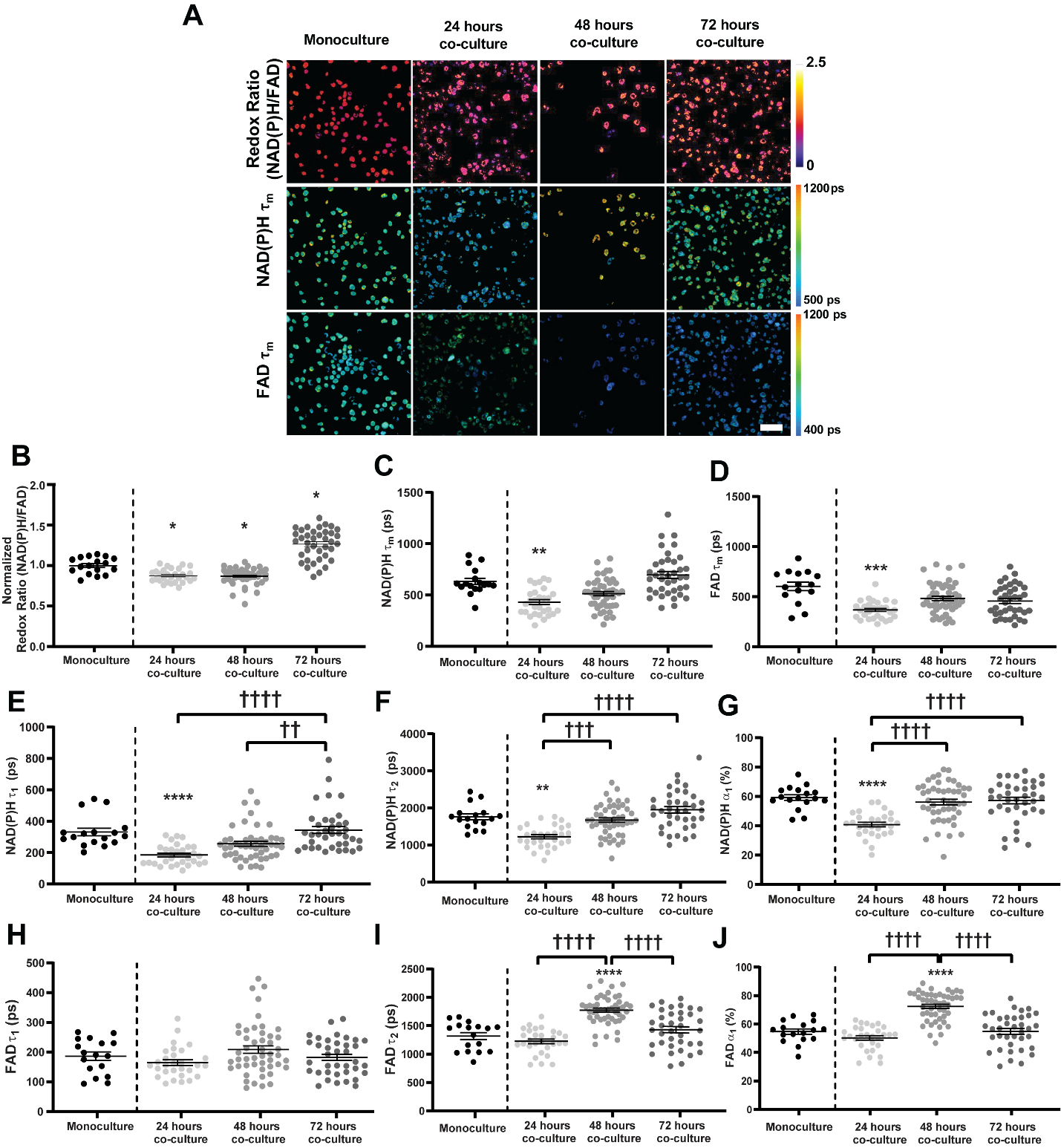

**Supplementary Figure 5: Metabolic changes in human THP-1s following co-culture with MDA-MB-231 human breast carcinoma.** A) Representative autofluorescence images demonstrate qualitative differences in the optical redox ratio, NAD(P)H τ_m_, and FAD τ_m_ in 3D monocultures of human THP-1s or co-cultures of MDA-MB-231 breast cancer and THP-1s. Scale bar = 50 µm. Quantitative trends in B) redox ratio, C) NAD(P)H τ_m_, D) FAD τ_m_, E) NAD(P)H τ_1_, F) NAD(P)H τ_2_, G) NAD(P)H α_1_, H) FAD τ_1_, I) FAD τ_2_, and J) FAD α_1_ of monocultures or co-cultures over 24, 48, and 72 hours;*,**,*** p<0.05,0.01,0.001 vs monoculture.

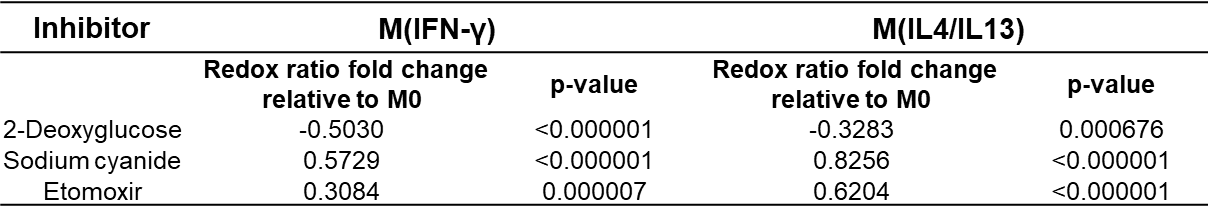

**Supplementary Table 1: Significance of redox ratio fold change in inhibitor-treated 2D cytokine-stimulated mouse macrophages**

**Supplementary Table 2: Mouse and Human Macrophage Polarization Gene Panels**

| **Gene Symbol** | **Refseq #** | | **Official Full Name** | **Associated Phenotype^76^** |
| --- | --- | --- | --- | --- |
| **Mouse gene panel** | | | | |
| Stat3 | NM_011486 | | signal transducer and activator of transcription 3 | M2-like |
| Vegfa | NM_001025 | | vascular endothelial growth factor A | M2-like |
| Ccl5 | NM_013653 | | chemokine (C-C motif) ligand 5 | M1-like |
| Il10 | NM_010548 | | interleukin 10 | M2-like |
| Ptgs2 | NM_011198 | | prostaglandin-endoperoxide synthase 2 | M2-like |
| Ccl2 | NM_011333 | | chemokine (C-C motif) ligand 2 | M1-like |
| Il23a | NM_031252 | | interleukin 23, alpha subunit p19 | M1-like |
| Ccl22 | NM_009137 | | chemokine (C-C motif) ligand 22 | Mixed |
| Il12b | NM_008352, XM_006532306 | | interleukin 12b | Mixed |
| Il1b | NM_008361, XM_006498795 | | interleukin 1 beta | M1-like |
| Il6 | NM_031168 | | interleukin 6 | Mixed |
| Nos2 | NM_010927, XM_006532446 | | nitric oxide synthase 2, inducible | Mixed |
| Cxcl5 | NM_009141 | | chemokine (C-X-C motif) ligand 5 | M1-like |
| Tnf | NM_013693, NM_001278601 | | tumor necrosis factor | Mixed |
| Hsp90ab1 | NM_008302 | | heat shock protein 90 alpha (cytosolic), class B member 1 | Reference gene |
| Pgk1 | NM_008828 | | phosphoglycerate kinase 1 | Reference gene |
| **Human gene panel** | | | | |
| Vegfa | | NM001025366 | vascular endothelial growth factor A | M2-like |
| Il10 | | NM_000572 | interleukin 10 | Mixed |
| Ptgs2 | | NM_000963 | prostaglandin-endoperoxide synthase 2 | M2-like |
| Ccl2 | | NM_002982 | chemokine (C-C motif) ligand 2 | M1-like |
| Ccl22 | | NM_00990 | chemokine (C-C motif) ligand 22 | M2-like |
| Il12b | | NM _00187 | interleukin 12b | Mixed |
| Il1b | | NM_000576 | interleukin 1 beta | M1-like |
| Il6 | | NM_000600 | interleukin 6 | Mixed |
| Nos2 | | NM_000625 | nitric oxide synthase 2, inducible | Mixed |
| Tnf | | NM_000594 | tumor necrosis factor | M1-like |
| Csf1 | | NM_000757 | colony stimulating factor 1 (macrophage) | Mixed |
| Tgfb1 | | NM_000660 | transforming growth factor, beta 1 | Mixed |
| Actb | | NM_001101 | actin, beta | Reference gene |
| B2m | | NM_004048 | beta-2-microglobulin | Reference gene |

**Supplementary Table 3: Coefficients of variation (CV) and significance of CV equality for optical redox ratio in 2D cytokine-stimulated mouse macrophages and 3D mouse monocultures and co-cultures**

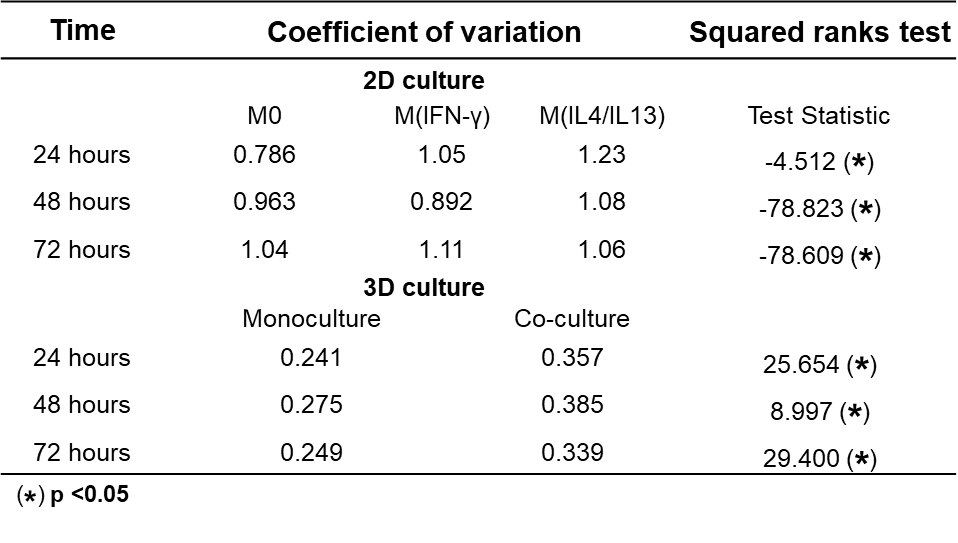
